# Serpine1 plays a critical role in Th17 cell differentiation and psoriasis pathogenesis

**DOI:** 10.1101/2025.05.08.652800

**Authors:** Kaidireya Saimaier, Ling Xie, Chun Wang, Jie Lv, Guangyu Liu, Sanxing Han, Chunliang Li, Changsheng Du

## Abstract

A large body of evidence indicates that Th17 cells play essential roles in mucosal immune responses and trigger autoimmune diseases, including multiple sclerosis, inflammatory bowel disease, and psoriasis. Targeting Th17 cells holds promise for therapeutic innovation. However, the molecular mechanisms underlying Th17 cell differentiation remain poorly understood. Here, we found that Serpine1 was preferentially induced in Th17 cells. Genetic ablation of Serpine1 inhibited Th17 cell polarization *in vitro*, reproducing the phenotype caused by pharmaceutical inhibition against Serpine1 product PAI-1. Moreover, we confirmed that, despite normal T-cell initiation *in vivo*, deletion or inhibition of Serpine1 significantly attenuated disease severity and notably reduced skin inflammation in an IMQ-induced psoriasis model. Notably, this protective effect was also recapitulated in the CD4 conditional knockout mouse pathogenic model, ruling out the non-T cell autologous effect of Serpine1. Collectively, our results revealed a critical role of Serpine1 in regulating Th17 cell differentiation and autoimmune diseases, suggesting that Serpine1 is a promising therapeutic target for autoimmune disease treatment.

**Significance:** Using multiple genetic and pharmaceutical ablation model systems, we have demonstrated that Serpine1 plays a critical role in regulating Th17 differentiation and psoriasis pathogenesis. Serpine1 intervention attenuates disease symptoms in a psoriatic mouse model in a Th17 cell-dependent manner. This finding extends our understanding of Serpine1’s role in inflammatory diseases, highlighting its therapeutic potential.

## Introduction

Psoriasis is a common immune-mediated skin disease characterized by infiltration of inflammatory cells and hyperproliferation of epidermal keratinocytes (1, 2). It is estimated that approximately 125 million people worldwide now suffer from psoriasis, leading to a heavy burden on individuals and public health systems (3, 4). While the exact pathogenesis has not been completely elucidated, established work has demonstrated Th17 cells as one of the key drivers in the progression of psoriasis (5–7). Genome-wide association studies (GWAS) have confirmed that multiple psoriasis susceptibility loci are closely associated with dysregulated Th17 cell differentiation (7–9). Clinical observations have also demonstrated that Th17 cell populations and corresponding cytokine levels are elevated at lesion sites and in the peripheral blood of psoriasis patients (10–12). Genetic studies confirm that knockdown of the characteristic Th17 cell transcription factor RORγt completely inhibits psoriasis-like skin inflammation in mice (13). Further mechanistic studies revealed that cytokines such as IL-17A and IL-22 secreted by Th17 cells induce rapid proliferation and impaired differentiation of keratinocytes in psoriasis and exacerbate the inflammatory cascade at the lesion site (14, 15). Therefore, understanding the molecular mechanisms underlying Th17 cell regulation and the development of therapeutic strategies targeting Th17 cell function is crucial for treating psoriasis.

Th17 cells are a subset of CD4^+^ T cells induced by IL-6 and TGF-β, specifically expressing IL-17A/F and transcription factors RORγt/RORα (16–20). Homeostatic Th17 cells protect the mucosal barrier from extracellular bacteria and fungi infections, whereas pathogenic Th17 cells induced by IL-23 and IL-1b trigger autoimmune diseases like psoriasis (21–24). Targeting approaches against Th17-related cytokines (TNF-α, IL-23, and IL-17A) have been approved by the US Food and Drug Administration (FDA) for clinical use, significantly enhancing the therapeutic efficacy of psoriasis (25–27). However, challenges remain due to variable therapeutic responses, risks of drug tolerance, and infection-related adverse events (28, 29). These limitations indicate an insufficient understanding of the plasticity, tissue-specific activation, and other regulatory mechanisms governing Th17 cells. Therefore, there is an urgent need for a comprehensive exploration of Th17 regulatory mechanisms to advance novel therapeutic strategies for autoimmune diseases.

Th17 cell functions depend on their secreted cytokine network. Developments of therapeutic targets for Th17 cells have long focused on classical inflammatory cytokines. In contrast, the potential regulatory roles of other secreted proteins in Th17 cell differentiation and autoimmune pathology remain poorly understood. As one of the most abundant types of proteins, secreted proteins make up about one-tenth of the genome and are involved in various biological processes such as signaling pathways, blood coagulation, and immune defense (30). Secreted proteins exist in free form in bodily fluids such as blood and urine, making them ideal non-invasive biomarkers for the dynamic monitoring of disease progression and therapeutic response (31). Additionally, the extracellular distribution of secreted proteins renders them effective targets for intervention by antibody drugs or small molecule compounds (30, 32). Recent studies have shown that diverse “non-classical” secreted proteins, including growth factors, matrix metalloproteinases, and extracellular vesicles, may influence Th17 cell function and autoimmune disease pathogenesis (33–35). Systematic identification and functional validation of these factors could enable novel multi-target therapies, addressing the current limitations of cytokine-focused treatments.

Here, we reported that Serpine1 is a Th17-specific secreted factor during T-cell differentiation. Systematic genetic ablation and pharmacological inhibition of Serpine1 impaired the polarization of Th17 cells. Moreover, Serpine1 deficiency reduced inflammation and attenuated disease severity in an IMQ-induced psoriasis mouse model. Consistent with findings in *de novo* knockouts, CD4-specific Serpine1 knockout mice exhibited comparable protection against psoriasis, demonstrating that Serpine1 modulates psoriatic pathology by regulating Th17 cell differentiation. Therapeutic targeting of PAI-1, the bioactive product of Serpine1, likewise attenuated disease symptoms and reduced skin thickness in a psoriasis mouse model. Given the importance of IL-17 signaling in psoriasis, these findings would provide new insights into the function of Serpine1 as a potential diagnostic factor and therapeutic target.

## Results

### Serpine1 is specifically expressed in Th17 cells

Naive CD4^+^ T cells can differentiate into distinct subpopulations (Th0, Th1, Th17, and Treg) *in vitro* in response to various stimulation conditions (Fig. 1A). By characterizing transcription profiling in different Th cell subpopulations and further evaluating the targeting possibility, we focused on the Serpine1 gene, which was redundantly expressed in Th17 cells. Serpine1 encodes plasminogen activator inhibitor-1 (PAI-1), which exerts antifibrinolytic effects by binding and inactivating tissue-type plasminogen activator (tPA) and urokinase-type plasminogen activator (uPA) (36, 37). However, other than selective expression, it is unclear whether Serpine1 functionally engages in the regulation of Th17 cell differentiation.

**Fig. 1.**
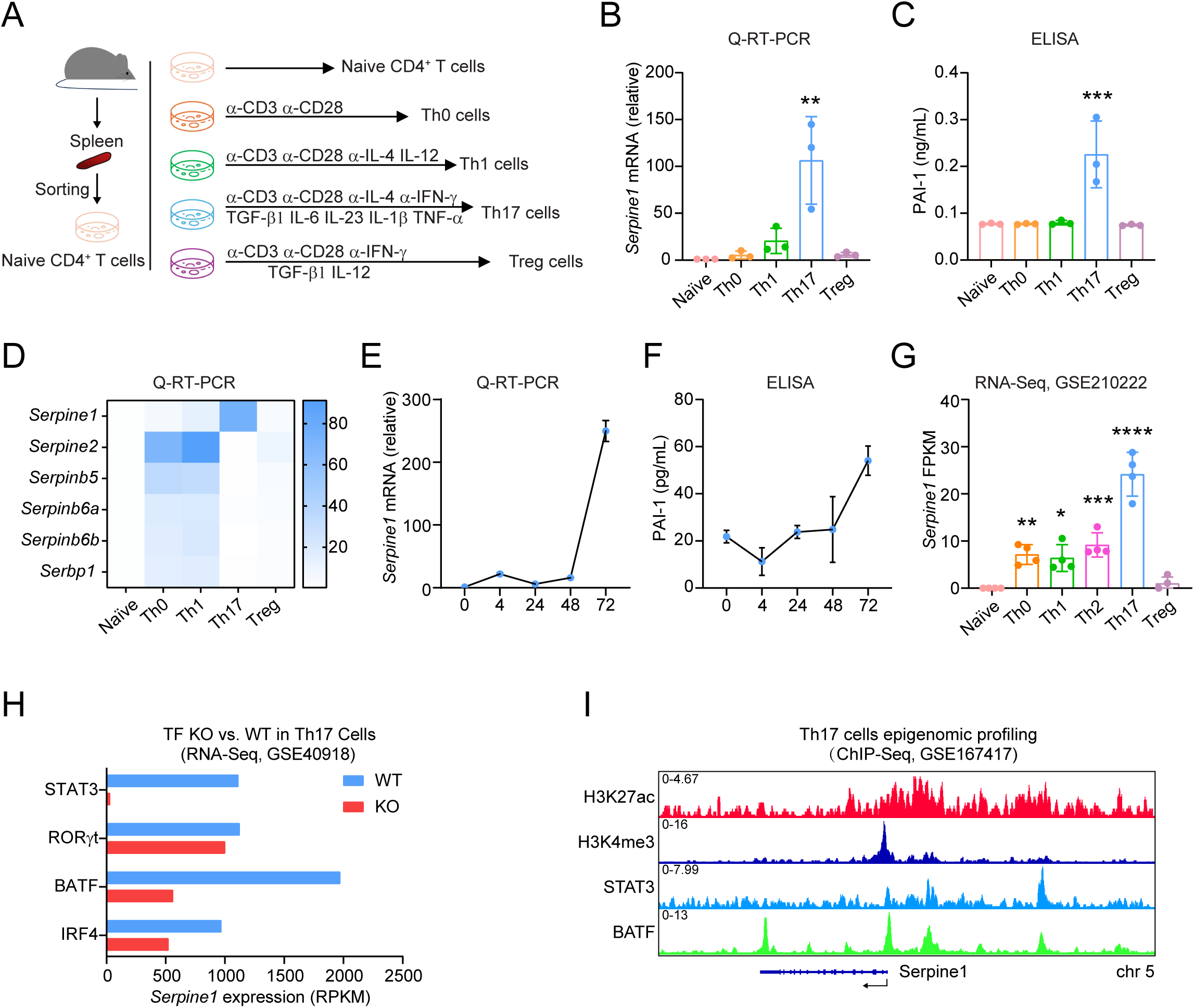
Th17 cells specifically express Serpine1. (A) Schematic diagram of *in vitro* differentiation of naive CD4^+^ T cells. (B) q-RT-PCR analysis of *Serpine1* mRNA levels in different Th cell subpopulations. (C) ELISA assay for PAI-1 expression levels in supernatants of different Th cell subpopulations. (D) qRT-PCR analysis of mRNA levels of other Serpin family members in Th17 cells. (E) qRT-PCR analysis of Serpine1 mRNA levels in Th17 cells at 0, 4, 24, 48, and 72h. (F) ELISA assay of PAI-1 expression levels in Th17 cell supernatants at 0, 4, 24, 48, and 72h. (G) Bar plot of RPKM expression values of Serpine1 transcripts in TF KO (Irf4^-/-^, Batf^-/-^, Rorc(t)^GFP/GFP^, Stat3^fl/fl^ CD4-Cre) versus WT 48-hour Th17 cell polarization culture. (H) ChIP-seq tracks for TFs and H3K27ac and H3K4me3 peaks enriched at the promoter of Serpine1 in 48-hour Th17 cell cultures. Data are presented as mean ± SEM. Statistical analysis was carried out by one-way ANOVA (B, C, and G). \**P*<0.05, \*\**P*<0.01, \*\*\**P*<0.001, \*\*\*\**P*<0.0001.

To investigate the specific expression pattern of Serpine1 in Th17 cells, we assessed its mRNA levels across different Th cell subsets using quantitative RT-PCR. We found that Serpine1 mRNA was relatively higher in Th17 cells compared to other subpopulations, consistent with our transcriptome profiling (Fig. 1B). Serpine1 encodes the PAI-1 protein, which is secreted as an active form into the extracellular environment (38). Consequently, we measured PAI-1 levels in the supernatants of Th cells during *in vitro* differentiation. We observed a significant increase in PAI-1 in the supernatant of Th17 cells (Fig. 1C). PAI-1 is a prominent member of the serine protease inhibitor (serpin) superfamily E. Despite the high degree of structural similarity among serpins, their functional roles are remarkably diverse (39). We also examined other members of the serine protease inhibitor family and found that only PAI-1 was selectively expressed in Th17 cells (Fig. 1D). Time-course analysis of Serpine1 transcription during Th17 cell differentiation revealed that Serpine1 is predominantly expressed at the late stages of polarization, along with its protein product, PAI-1 (Fig. 1E and 1F). These findings align with the publicly available RNA sequencing dataset (GSE210222), which indicates that Serpine1 mRNA is significantly elevated in Th17 cells (Fig. 1G).

To investigate the potential regulatory basis of Th17 cell-specific Serpine1 expression, we assessed the transcriptomic changes and binding occupancy of Th17 cell-related transcription factors (TFs), including IRF4, BATF, STAT3, and RORγt (40). Compared with wild-type (WT) settings, deletion of BATF, IRF4, and STAT3 notably reduced Serpine1 mRNA expression in Th17 cells (Fig. 1H). Interestingly, the Th17 cell lineage-defining transcription factor RORγt did not affect Serpine1 expression. In addition, we revisited the chromatin immunoprecipitation sequencing (ChIP-seq) data (GSE167417). We found that BATF and STAT3 occupy the promoter and upstream regulatory elements of Serpine1, which were marked by H3K27ac and H3K4me3 modifications (Fig. 1I). Collectively, these findings indicate that Serpine1 is specifically expressed in Th17 cells and is likely regulated by a combination of the pioneer transcription factors.

### Serpine1 promotes Th17 cell polarization

Given the specific expression of Serpine1 in Th17 cells, we hypothesized that Serpine1 likely plays a critical role in regulating Th17 cell differentiation. We employed Serpine1 knockout (Serpine1^-/-^) mice for *ex vivo* and *in vivo* functional characterization of Th17 cells. The genetic model was established through homologous recombination, replacing the Serpine1 promoter and exons 1-9 with the neomycin resistance cassette (Fig. 2A and SI Appendix, Fig. S1A) (41). Complete knockout in Serpine1^-/-^ mice was confirmed by genotyping PCR (SI Appendix, Fig. S1B). To assess deletion efficiency, we examined PAI-1 levels in the serum of Serpine1 knockout mice. Enzyme-linked immunosorbent assays (ELISA) showed that PAI-1 expression was barely detectable in Serpine1^-/-^ mice serum compared to WT controls (SI Appendix, Fig. S1C).

**Fig. 2.**
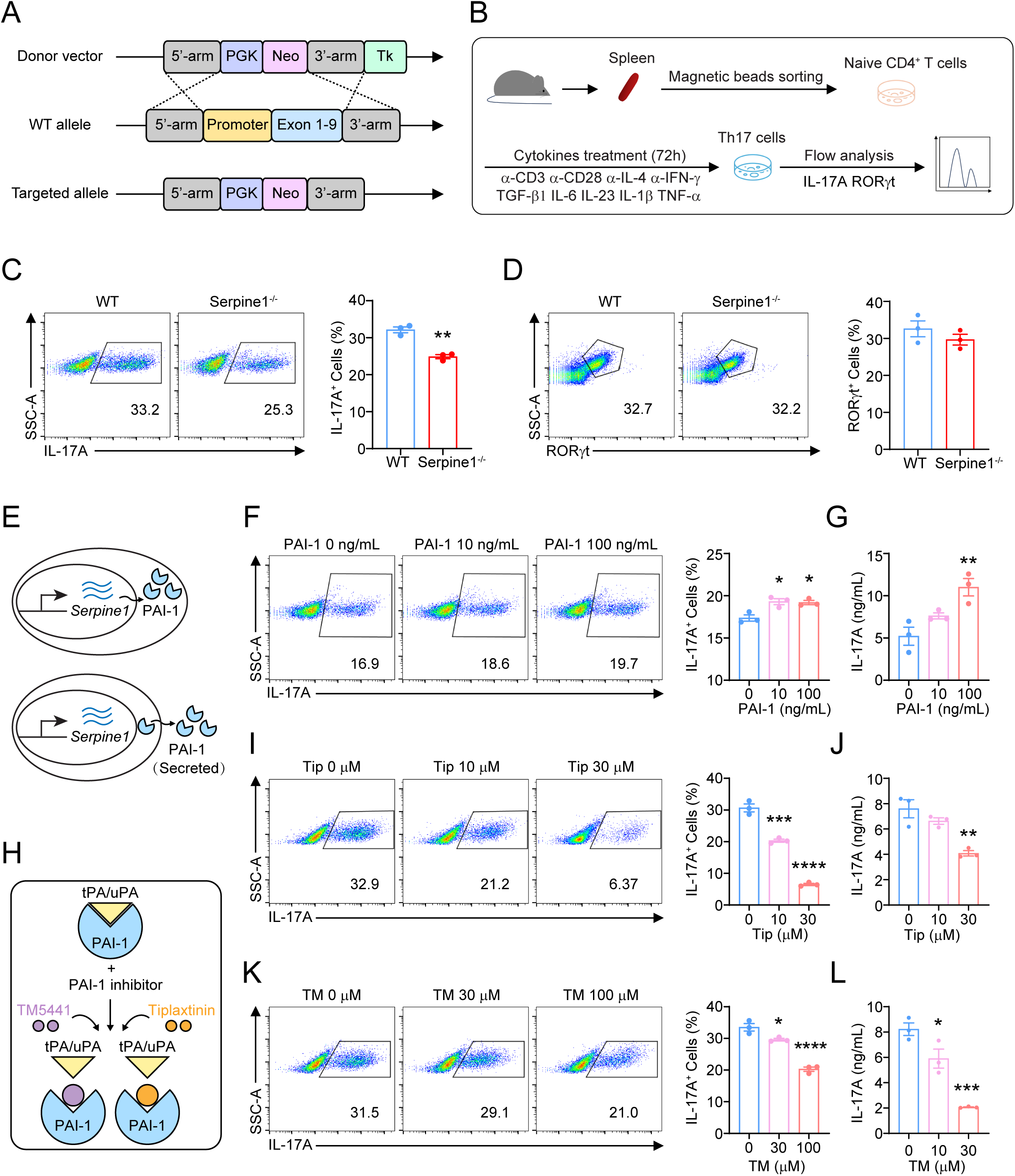
Serpine1 promotes Th17 cell differentiation. (A) Schematic diagram of the strategy to knock out PAI-1. (B) Schematic diagram of Th17 cell differentiation *in vitro*. (C) Flow cytometry detection of Serpine1^-/-^ and WT Th17 cells (IL-17A^+^) differentiated *in vitro*. Representative flow cytometry images (left). IL-17A^+^ cell frequency (right). (D) Flow cytometry detection of the transcription factor RORγt in differentiated Serpine1^-/-^ and WT Th17 cells *in vitro*. Representative flow cytometry images (left). RORγt^+^ cell frequency (right). (E) Schematic diagram of PAI-1 protein expression and secretion. (F) Flow cytometry detection of *in vitro* differentiated Th17 cells in the presence or absence of PAI-1 recombinant protein. Representative flow cytometry images (left). IL-17A^+^ cell frequency (right). (G) ELISA assay of IL-17A expression in supernatants of *in vitro* differentiated Th17 cells in the presence or absence of PAI-1 recombinant protein. (H) Schematic diagram of the mechanisms of the PAI-1 inhibitors Tiplaxtinin and TM5441. (I) Flow cytometry detection of *in vitro* differentiated Th17 cells in the presence or absence of Tiplaxtinin. Representative flow cytometry images (left). IL-17A^+^ cell frequency (right). (J) ELISA assay of IL-17A expression in supernatants of *in vitro* differentiated Th17 cells in the presence or absence of Tiplaxtinin. (K) Flow cytometry detection of *in vitro* differentiated Th17 cells in the presence or absence of TM5441. Representative flow cytometry images (left). IL-17A^+^ cell frequency (right). (L) ELISA assay of IL-17A expression in supernatants of *in vitro* differentiated Th17 cells in the presence or absence of TM5441. Data are presented as mean ± SEM. Statistical analysis was carried out by unpaired Student’s t-test (C and D) or one-way ANOVA test (F-L). \**P*<0.05, \*\**P*<0.01, \*\*\**P*<0.001, \*\*\*\**P*<0.0001.

Subsequently, we examined the effect of Serpine1 deletion on the differentiation of Th17 cells through the *in vitro* differentiation of CD4^+^ T cells. Naive CD4^+^ T cells from WT and Serpine1 knockout mice were isolated and cultured under Th17 cell polarization conditions for 72 h. The impact of Serpine1 deletion on Th17 cell differentiation *in vitro* was then assessed using flow cytometry (Fig. 2B). We found that the absence of Serpine1 significantly impaired Th17 cell differentiation compared to WT controls (Fig. 2C). However, Serpine1 did not seem to affect RORγt expression, indicating that Serpine1 likely promotes Th17 cell differentiation in a RORγt-independent manner (Fig. 2D). Moreover, with the corresponding *in vitro* differentiation conditions, we also examined the effect of Serpine1 on the differentiation of other Th lineages (SI Appendix, Fig. S1D). We noticed a normal frequency of Serpine1^-/-^ Treg cells but a reduced frequency of Th1 cells compared with WT controls (SI Appendix, Fig.S1E and F).

Serpine1 encodes the PAI-1 protein and is then secreted extracellularly (Fig. 2E). To investigate whether exogenous PAI-1 modulates Th17 differentiation, we supplemented cultures with recombinant PAI-1. The addition of PAI-1 recombinant protein promoted Th17 cell differentiation and increased IL-17A secretion in the supernatant (Fig. 2F and G). Tiplaxtinin (Tip) and TM5441 (TM) are the most commonly used PAI-1 inhibitors, which potently block PAI-1 function by preventing the formation of a stable covalent complex between PAI-1 and the substrate (Fig. 2H; SI Appendix, Fig. S1G and H) (42, 43). Next, we assessed the pharmacological effects of these PAI-1 inhibitors on Th17 cell differentiation *in vitro*. Th17 cells were treated with either the PAI-1 inhibitors or DMSO as a control for 72 hours. Flow cytometry analysis showed that Tip inhibited the differentiation of Th17 cells in a dose-dependent manner (Fig. 3I). At the same time, a corresponding reduction in IL-17A expression was observed in the supernatant of Th17 cells (Fig. 2J). Notably, this inhibitory effect was recapitulated by another PAI-1 inhibitor, TM (Fig. 2K and L). Meanwhile, we examined the effect of PAI-1 inhibitors on other Th lineages. We found that only high concentrations of PAI-1 inhibitors reduced Th1 cell differentiation, suggesting that Th17 cells are more sensitive to PAI-1 inhibitors (SI Appendix, Fig. S1I and J). Consistent with the results of genetic intervention, PAI-1 inhibitors did not affect Treg cell differentiation (SI Appendix, Fig. S1I and J). In conclusion, our findings demonstrate that Serpine1 promotes differentiation of Th17 and Th1 cells, not Treg cells.

**Fig. 3.**
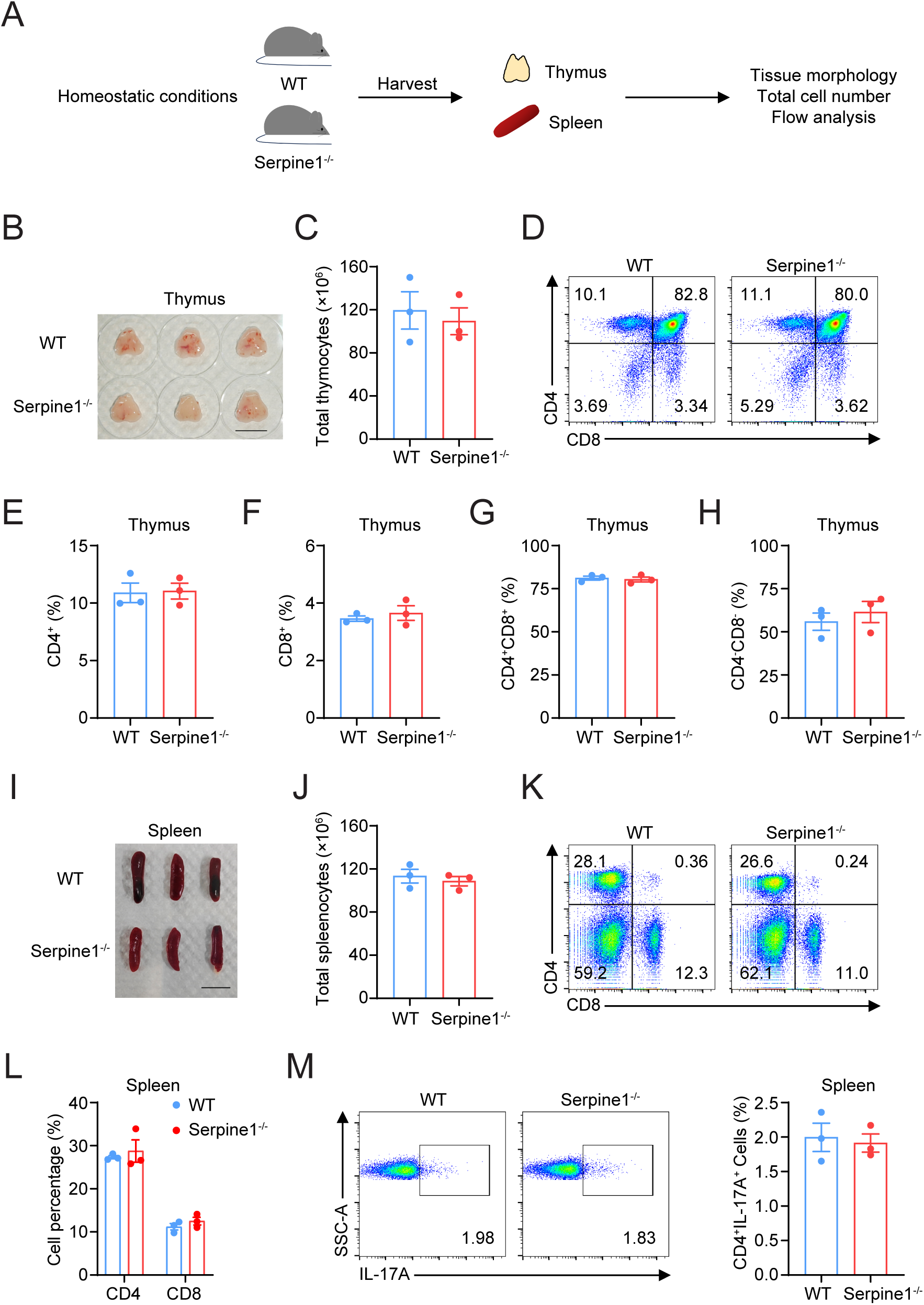
Serpine1 is not essential for homeostatic T cells. (A) Schematic diagram of the experimental design for analyzing immune cells in the thymus and spleen of Serpine1^-/-^ and WT mice at steady state. (B) Pictures of the thymus morphology of Serpine1^-/-^ and WT mice. (C) Total thymocyte counts of Serpine1^-/-^ and WT mice. (D) Representative flow analysis plots of CD4^+^ and CD8^+^ in Serpine1^-/-^ and WT thymocytes. (E-H) Flow cytometry detection for (E) CD4^+^, (F) CD8^+^, (G) CD4^+^CD8^+^, and (H) CD4^-^CD8^-^ cell ratios in Serpine1^-/-^ and WT thymocytes. (I) Pictures of spleen morphology of Serpine1^-/-^ and WT mice. (J) Total Splenocyte counts of Serpine1^-/-^ and WT mice. (K) Representative flow analysis plots of CD4^+^ and CD8^+^ in Serpine1^-/-^ and WT Splenocytes. (L) Flow assay for CD4^+^ and CD8^+^ cell ratios in Serpine1^-/-^ and WT Splenocytes. (M) Flow cytometry detection of the proportion of Th17 cells (CD4^+^IL-17A^+^) in the spleens of Serpine1^-/-^ and WT mice. Representative flow cytometry images (left). Cell frequencies (right). Data are presented as mean ± SEM. Statistical analysis was carried out using an unpaired Student’s t-test (E-H and L-M).

### Serpine1 deficiency does not affect peripheral T cell homeostasis

To further consolidate the specific role of Serpine1 in pathogenesis, we also examined the effect of Serpine1 deletion on normal T-cell development under steady-state conditions. Thymus and spleen tissues were isolated from WT and Serpine1^-/-^ mice at homeostasis. We assessed the effects of Serpine1 on peripheral immune organs through morphology evaluation, cell counting, and flow cytometry analysis (Fig. 3A). Compared with WT mice, Serpine1 deletion did not affect thymus morphology or total cell number in mice (Fig. 3B and C). We further examined the distribution of the four major thymic populations. We found that the percentage of CD4^-^ CD8^-^ double-negative (DN), CD4^+^CD8^+^ double-positive (DP), and CD4^+^ and CD8^+^ single-positive (SP) thymocytes in Serpine1^-/-^ mice was similar to those in WT controls (Fig. 3D-H). These results suggest that CD4^+^ T cell development in the thymus is not affected by Serpine1 deletion.

Next, we investigated the effects of Serpine1 deletion on the spleen. Serpine1 deletion did not affect spleen morphology or total cell count (Fig. 3I and J). Analysis of the frequencies of CD4^+^ T cells and Th17 cell subsets in the spleens of mice showed no significant differences between Serpine1^-/-^ and WT mice. (Fig. 3K-M). In addition, we examined whether Serpine1 deletion affected the frequency of CD4^+^ T cells and Th17 cells in secondary lymphoid organs (SI Appendix, Fig. S2A). As a result, the proportions of CD4^+^ T cells and Th17 cells in mesenteric lymph nodes (MLN) and draining lymph nodes (DLN) were also similar between WT and Serpine1^-/-^ mice (SI Appendix, Fig. S2B-C). Th17 cells play pathogenic or protective functions in the liver and lung (44, 45). Therefore, we also isolated Th17 cells in the liver and lung to examine whether Serpine1 deletion affects the proportion of T cells in different organs (SI Appendix, Fig. S2A). Flow cytometry showed that Serpine1 deletion did not affect the ratio of CD4^+^ T cells or Th17 cells in the liver and lung at homeostasis (SI Appendix, Fig. S2D and E). Collectively, these results suggest that Serpine1 does not affect thymic T cell development and peripheral T cell homeostasis.

### Serpine1 deficiency alleviates psoriasis pathology

Given the essential role of Serpine1 in inducing Th17 cell differentiation *in vitro*, we sought to evaluate its pathophysiological relevance in Th17-mediated autoimmunity by an *in vivo* approach. Psoriasis is an autoimmune disease driven predominantly by Th17 cells (46). IMQ-induced skin inflammation in mice is a psoriasis-like model that can further elucidate the pathogenic mechanisms of psoriasis and evaluate new therapies (47). Therefore, we applied IMQ cream on the backs of mice for seven consecutive days, recorded disease scores daily, and collected mouse skin, serum, and spleen on day 7 for subsequent analysis (Fig. 4A). As expected, IMQ-treated mice showed significant psoriasis-like lesions compared to the Vaseline-treated group (Fig. 4B-E). Notably, we observed a substantial increase in PAI-1 expression in the serum of IMQ-treated mice (SI Appendix, Fig. S3A). Likewise, mRNA levels of Serpine1 in the skin and spleen were elevated, indicating that Serpine1 may play a functional role in the psoriasis disease process (SI Appendix, Fig. S3B and C).

**Fig. 4.**
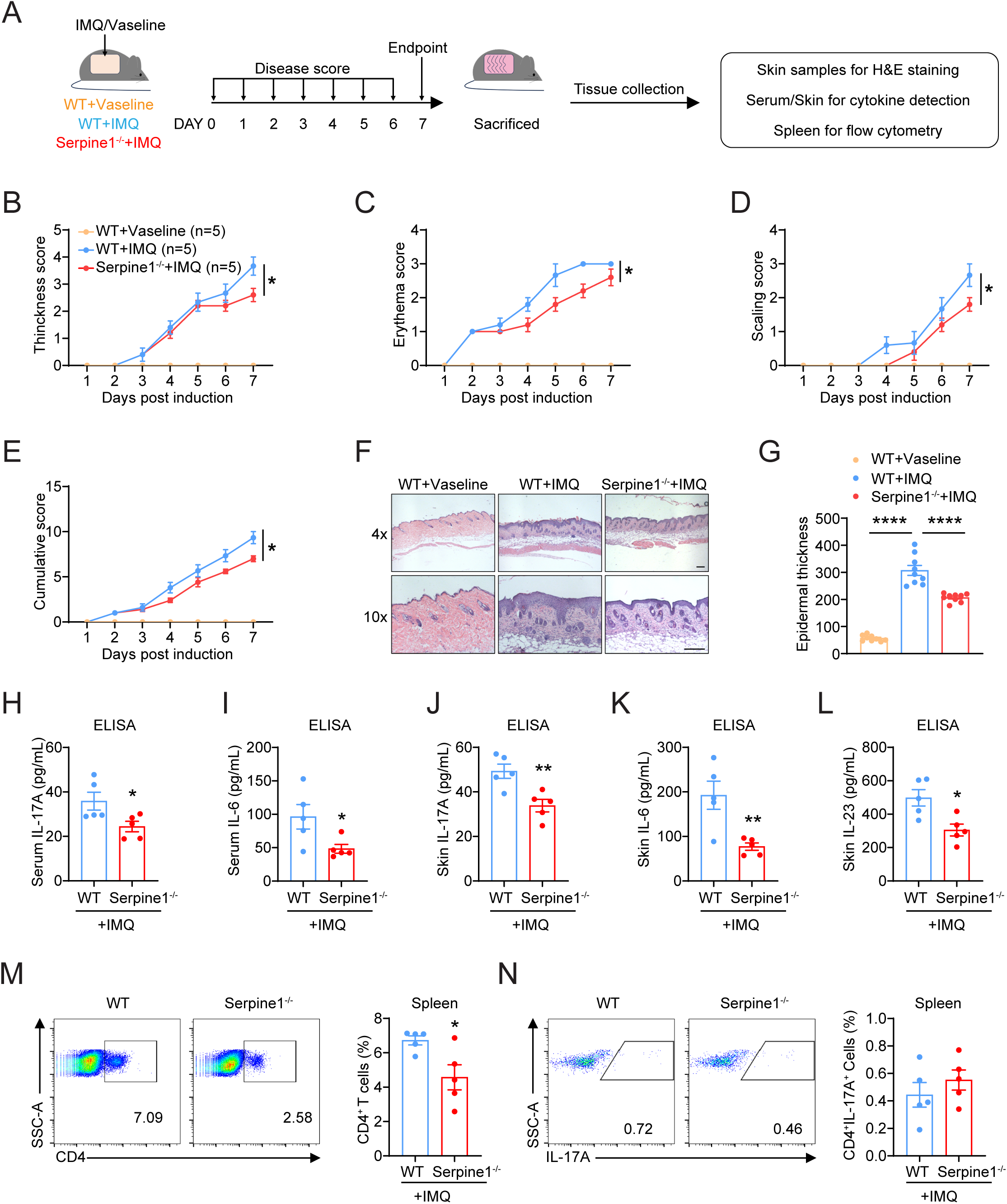
Serpine1 deficiency attenuates psoriasis symptoms in mice. (A) Schematic diagram of psoriasis mouse modeling and experimental design. (B-E) Scoring curves of back skin (B) thickness, (C) scaling, (D) erythema, and (E) Cumulative score. (F) H&E staining of skin sections. Scale bar=100 μm. (G) The epidermal thickness of mouse H&E-stained sections was analyzed by Image Pro. (H-I) Detection of (H) IL-17A and (I) IL-6 expression levels in mouse serum by ELISA. (J-L) Detection of (J) IL-17A, (K) IL-6, and (L) IL-23 expression Levels in Mouse Skin Homogenates by ELISA. (M) Flow cytometry detection of the CD4 cells (CD4^+^) ratio in the Spleen. Representative flow cytometry images (left). Cell frequencies (right). (N) Flow cytometry detection of the ratio of Th17 cells (CD4^+^IL-17A^+^) in the Spleen. Representative flow cytometry images (left). Cell frequencies (right). Data are presented as mean ± SEM. Statistical analysis was carried out by a two-way ANOVA test (B-E), one-way ANOVA test (G), or unpaired Student’s t-test (H-N). \**P*<0.05, \*\**P*<0.01, \*\*\*\**P*<0.0001.

By measuring skin thickness, erythema, and scaling, we found that Serpine1^-/-^ mice exhibited reduced disease severity compared to WT controls (Fig. 4B-E). Hematoxylin and eosin (H&E) staining of skin tissues showed that after IMQ-induced psoriasis development, Serpine1^-/-^ mice had reduced epidermal thickness compared to control mice (Fig. 4F and G). IL-17A, IL-6, and IL-23 are cytokines closely associated with the development of psoriatic lesions, leading to aberrant keratinocyte differentiation and increased skin inflammation (48). We also examined the secretion of inflammatory cytokines in serum and skin of IMQ-induced psoriasis mice. Compared to WT controls, Serpine1^-/-^ mice showed significantly reduced IL-17A and IL-6 levels in the serum (Fig. 4H and I) and IL-17A, IL-6, and IL-23 levels in the skin (Fig. 4J-L).

Th17 cell differentiation is initiated in the spleen and peripheral lymph nodes, where dendritic cells (DCs) present antigens to naïve CD4^+^ T cells and drive Th17 cell differentiation (49, 50). Mature Th17 cells subsequently upregulate the chemokine receptor CCR6, acquiring the skin-homing capacity to infiltrate inflamed tissues (49). To further investigate the role of Serpine1 in Th17 cell-mediated autoimmune diseases, we analyzed the ratio of CD4^+^ T cells and Th17 cells in the spleen (Fig. 4A) and lymph nodes (SI Appendix, Fig. S3D) using flow cytometry. We found that CD4^+^ T cells in splenocytes were reduced in Serpine1^-/-^ psoriatic mice, while Th17 cells remained unchanged (Fig. 4M and N). Moreover, flow cytometry analysis of the DLN and MLN revealed the decreased Th17 cells in the Serpine1^-/-^ mice (SI Appendix, Fig. S3E and F). These findings suggest that Serpine1 deletion alleviated IMQ-induced psoriasis disease symptoms and reduced systemic and focal inflammation.

### Deletion of Serpine1 in CD4^+^ T cells attenuates psoriasis

Given the constitutive expression of Serpine1 across multiple tissue types, conventional whole-body knockout (KO) models may not definitively attribute observed phenotypes to Th17 cells or T lymphocytes (51). To mitigate this limitation, we employed a conditional knockout (cKO) strategy utilizing the Cre-loxP system to achieve T cell-specific deletion of Serpine1. This approach enables precise genetic ablation of Serpine1 in CD4^+^ T cells through Cre recombinase activity driven by the CD4 promoter (SI Appendix, Fig. S4A and B). Therefore, we can define T cell-autonomous functions while preserving systemic Serpine1 expression in other cell populations. The qPCR and ELISA validated efficient Serpine1 ablation in CD4^+^ T cells isolated from Serpine1 cKO mice, showing that the coding region of Serpine1 was significantly shorter and the protein level of Serpine1 was markedly reduced (SI Appendix, Fig. S4C-E), confirming efficient T cell-restricted ablation. *In vitro* differentiation assays demonstrated that CD4^+^ T cell-specific ablation of Serpine1 impaired differentiation of Th17 and Th1 cells without affecting Treg cell differentiation (Fig. 5A and B; SI Appendix, Fig. S4F-H). Next, we subjected cKO mice to IMQ-induced psoriasis. Consistent with previous results in systemic knockout mice, CD4^+^ T cell-specific knockout of Serpine1 significantly attenuated the clinical severity of the disease, with significant reductions in thickness, erythema, and scaling scores (Fig. 5C-G). Histopathological analysis of H&E-stained lesional skin sections revealed that epidermal hyperplasia was significantly reduced in mice in the conditional knockout group compared to controls (Fig. 5H and I). Flow cytometric analysis of spleen and DLN cells revealed a marked decrease of CD4^+^ and IL-17A^+^ cell populations in cKO mice compared to controls (Fig. 5J and K; SI Appendix, Fig. S4I and J). A noticeable reduction in CD4^+^ T cells was observed in the MLN, but the proportion of IL-17A^+^ cells was unchanged (SI Appendix, Fig. S4K and L). Collectively, these data establish Serpine1 as a T cell-intrinsic regulator of Th17-driven psoriatic inflammation.

**Fig. 5.**
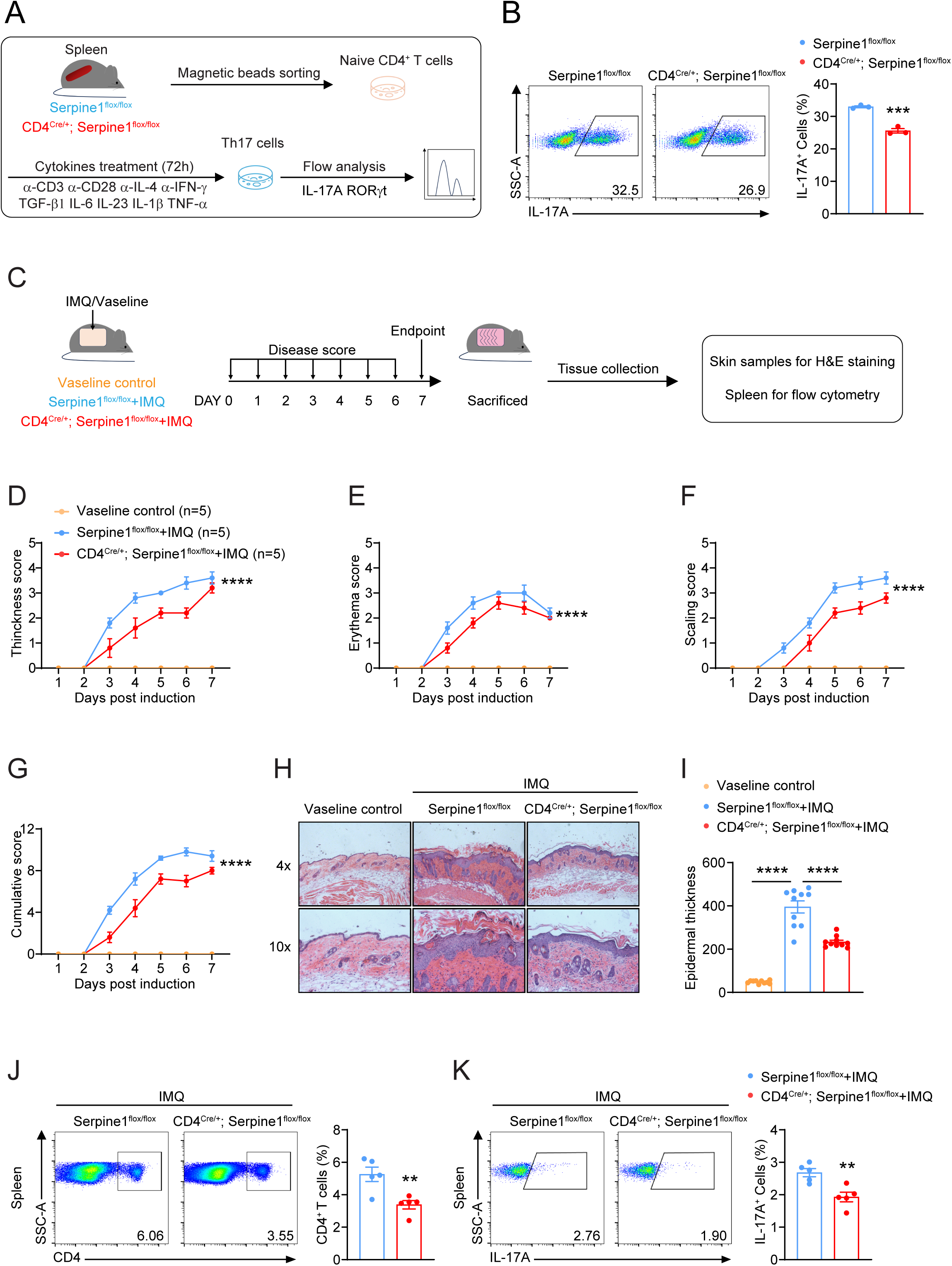
CD4 cell-specific knockdown of Serpine1 attenuates psoriasis symptoms in mice. (A) Schematic diagram of Th17 cell differentiation *in vitro*. (B) Flow cytometry detection of CD4^Cre/+^; Serpine1^flox/flox^ and Serpine1^flox/flox^ Th17 cells (IL-17A^+^) differentiated *in vitro*. Representative flow cytometry images (left). IL-17A^+^ cell frequency (right). (C) Schematic diagram of psoriasis mouse modeling and experimental design in CD4^+^-specific knockout Serpine1 mice. (D-G) Scoring curves of back skin (D) thickness, (E) scaling, (F) erythema, and (G) Cumulative score. (H) H&E staining of skin sections. Scale bar=100 μm. (I) The epidermal thickness of mouse H&E-stained sections was analyzed by Image Pro. (J) Flow cytometry detection of the CD4 cells (CD4^+^) ratio in the Spleen. Representative flow cytometry images (left). Cell frequencies (right). (K) Flow cytometry detection of the ratio of Th17 cells (CD4^+^IL-17A^+^) in the Spleen. Representative flow cytometry images (left). Cell frequencies (right). Data are presented as mean ± SEM. Statistical analysis was carried out by a one-way ANOVA test. \*\**P*<0.01, \*\*\**P*<0.01, \*\*\*\**P*<0.0001.

### PAI-1 inhibitor attenuates IMQ-induced psoriasis in mice

Previously, we observed that PAI-1 inhibitors Tip and TM inhibited Th17 cell differentiation *in vitro* (Fig. 2H-I). To explore interventions targeting Serpine1 for psoriasis treatment, the *in vivo* therapeutic efficacy of the Tip was assessed. In the IMQ-induced psoriasis-like mouse model, we administered Tip to mice for 7 days by gavage to test its inhibitory effect on disease progression (Fig. 6A). Histopathologic results of the skin showed that the PAI-1 inhibitor significantly reduced skin thickness compared to the control group (Fig. 6B and C). Additionally, we tested the effect of PAI-1 inhibitors on immune cells in the spleen of psoriatic mice. A decrease in CD4⁺ T cells was observed in the spleen (Fig. 6D). In contrast, the frequency of Th17 cells remained unchanged (Fig. 6D). Taken together, *in vivo,* inhibition of PAI-1 function can effectively alleviate the disease severity of psoriasis and provide a new strategy for treating psoriasis.

**Fig. 6.**
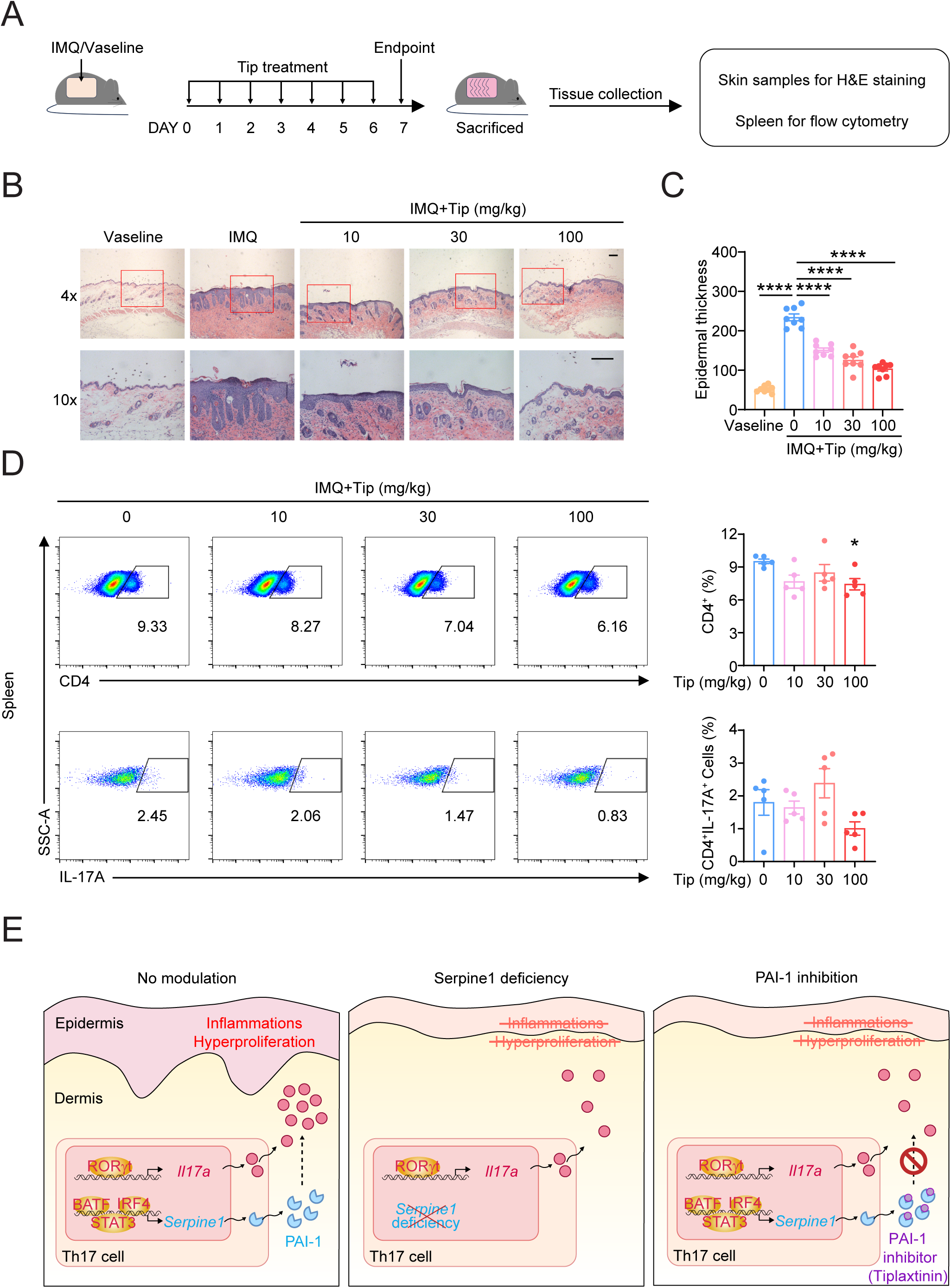
PAI-1 inhibitor Tiplaxtinin attenuates psoriasis in mice. (A) Mice were administered Tiplaxtinin by gavage for seven consecutive days, and IMQ was applied to induce psoriasis. Mice were sacrificed on the seventh day, and tissues were collected for subsequent analysis. (B) H&E staining of skin sections. Scale bar=100 μm. (C) The epidermal thickness of mouse H&E-stained sections was analyzed by Image Pro. (D) Flow cytometry detection of the ratio of CD4 cells (CD4^+^) to Th17 cells (CD4^+^IL-17A^+^) in mouse spleen. Representative flow cytometry images (left). Cell frequencies (right). (E) Schematic diagram summarizing this study. Our study revealed that Serpine1 promotes Th17 cell differentiation and is involved in Th17 cell-mediated autoimmune disease development. Serpine1 has therapeutic potential as a regulator of Th17 cells in treating autoimmune diseases. Data are presented as mean ± SEM. Statistical analysis was carried out by a one-way ANOVA test. \**P*<0.05, \*\*\*\**P*<0.0001.

In summary, we found that Th17 cell differentiation-associated transcription factors regulate Serpine1 and drive the specific expression of Serpine1 in Th17 cells. Functional annotation revealed that Serpine1 intrinsically promotes Th17 polarization during T-cell differentiation. Using a Th17-driven psoriasis model, both genetic ablation and pharmacological inhibition of Serpine1 attenuated disease pathology through selective suppression of Th17 differentiation. The above studies suggest that Serpine1 is a critical regulator of Th17 cells and is a potential target for autoimmune disease treatment (Fig. 6E).

## Discussion

While Th17 cells have been well established as central effectors in autoimmune disease pathogenesis, the lack of effective therapeutic targets among known regulators has impeded progress in treatment development. Here, we identified Serpine1 as a novel Th17 cell regulatory factor that has not been previously reported. We found that Serpine1 is highly expressed in Th17 cells and is regulated by Th17 cell-associated transcription factors. By establishing *in vitro* differentiation of CD4^+^ T cells, we found that genetic intervention or pharmacological inhibition of Serpine1 significantly impaired Th17 cell polarization. Under steady-state conditions, Serpine1 deficiency does not affect overall T cell development or peripheral Th17 cell distribution. Instead, in an IMQ-induced psoriasis model, both global and CD4 conditional knockout mice showed markedly attenuated skin inflammation, suggesting that Serpine1 affects the disease process by regulating Th17 cell differentiation. Finally, treatment with a PAI-1 inhibitor confirms the therapeutic potential of targeting Serpine1 in psoriasis. Together, our findings fill a critical knowledge gap in Th17 cell regulation and highlight Serpine1 as a promising therapeutic target for autoimmune diseases.

Previous studies of Serpine1 have focused on the function of the fibrinolytic system, but a growing body of research suggests that Serpine1 is associated with inflammation. Both Serpine1 and its protein product PAI-1 have been reported to be markedly upregulated in a range of autoimmune diseases, including multiple sclerosis, rheumatoid arthritis, and psoriasis, underscoring its potential involvement in chronic inflammatory pathologies (52–54). Genetic deficiency or pharmacological blockade of Serpine1 significantly reduced disease severity in MOG-induced EAE or DSS-induced colitis, indicating a pathogenic role of Serpine1 in inflammatory disease progression (55, 56). However, Serpine1’s function in immune cells, particularly T cells, remains unclear. Our study addresses this knowledge gap by identifying Serpine1 as a T cell-intrinsic regulator of Th17 differentiation. Similarly, systems biology analyses in IBD patient cohorts reveal that PAI-1 expression distinguishes patients with active disease and is directly associated with IL-17 signaling networks in inflamed mucosa (55). These data align to position Serpine1 as a critical nexus in Th17 differentiation and inflammation.

The upregulation of Serpine1 expression in Th17 cells is probably driven by multiple inflammatory environmental factors. Pro-inflammatory cytokine stimulation is a key initiator of T cell differentiation and autoimmune disease development. Previous studies have shown that TGF-β and IL-6 are necessary for T-cell differentiation (21). Intriguingly, Serpine1 expression has also been shown to be highly responsive to these cytokines (57, 58). The *Serpine1* mRNA level was induced more than 50-fold in TGF-β-treated human lung fibroblasts (57). In addition, it was found that endothelial cells induced PAI-1 expression through IL-6 trans-signaling (58). Consistent with these findings, we found that STAT3, a key transcription factor activated downstream of IL-6 receptor signaling, binds to regulatory regions of the Serpine1 gene and promotes its expression during Th17 cell differentiation. Despite these advances, the precise molecular mechanisms by which TGF-β and IL-6 collaboratively regulate Serpine1 expression remain to be fully elucidated. Future studies are necessary to further clarify the specific transcriptional regulatory mechanisms and upstream signaling pathways.

IL-17A/Th17 cells have emerged as key players in various inflammatory diseases, and they appear to be particularly prominent in skin inflammation (59). While the specific signaling pathways of PAI-1 in Th17 cell differentiation have not been well studied, existing evidence suggests that PAI-1 may regulate this process through several key pathways. PAI-1 is likely implicated in modulating MAPK pathways and NF-κB signaling, which are instrumental in mediating inflammatory responses and Th17 cell functions (60, 61). Future research involving systematic mechanistic studies and clinical sample validation is essential to clarify the detailed signaling networks associated with PAI-1. These insights could position PAI-1-related pathways as promising targets for treating autoimmune diseases, thereby facilitating the development of innovative and more effective therapeutic strategies.

Together, we have identified Serpine1 as a selective and critical determinant of Th17 cell differentiation and psoriasis. Hence, Serpine1 represents a potential target for therapeutic intervention in psoriasis.

## Methods

### Mice

C57BL/6J (Cat# N000013), B6.129S2-Serpine1^tm1Mlg/J^ (Cat# B000900), C57BL/6JGpt-Serpine1^em1Cflox^ /Gpt (T013287), and C57BL/6JGpt-Cd4^em1Cin(IRES-iCre)^/Gpt (T055135) mice were purchased from GemPharmatech Co., Ltd (Nanjing, China). Both genders were considered during the experiment. All mice were bred and maintained under a specific pathogen-free (SPF) facility. All experimental procedures were performed under approval by the Institutional Animal Care and Use Committee of Tongji University.

### CD4^+^ T cell purification and *in vitro* differentiation

Naïve CD4^+^ T cells are isolated from the spleens of 7-8 weeks old mice using Dynabeads® Untouched Mouse CD4 Cells Isolation Kit (Invitrogen, 11415D). Naïve CD4^+^ T cells are seeded in 96-well culture plates with 5*10^5^ cells/well and cultured in a complete RPMI 1640 medium (Gibco, C11875500BT) supplemented with 10% fetal bovine serum (ExCell Bio, FSD500), 50 μM β-ME (Gibco, 21985-023), and 100 U/mL penicillin/streptomycin (Biosharp, BL505A). The following cytokines and antibodies were added directly into the medium for differentiation of Th cell subpopulations: for Th0 cells: 2 μg/ml anti-CD3 antibody (Bio X Cell, BE0015-1), 2 μg/ml anti-CD28 antibody (Bio X Cell, BE0015-1); for Th1 cells: 2 μg/ml anti-CD3 antibody, 2 μg/ml anti-CD28 antibody, 10 ng/ml anti-IL-4 antibody (Bio X Cell, BE0045), 10 ng/ml IL-12 (PeproTech, 210-12); for Th17 cells: 2 μg/ml anti-CD3 antibody, 2 μg/ml anti-CD28 antibody, 10 μg/ml anti-IFN-γ antibody (Bio X Cell, BE0054), 10 μg/ml anti-IL-4 antibody, 3 ng/ml TGF-β (PeproTech, 100-21), 30 ng/ml IL-6 (PeproTech, 216-16), 10 ng/ml TNF-α (PeproTech, 211-11B), 10 ng/ml IL-23 (BioLegend, 589-002), 10 ng/ml IL-1β (PeproTech, 211-11B); for Treg cells: 2 μg/ml anti-CD3 antibody, 2 μg/ml anti-CD28 antibody, 10 μg/ml anti-IFN-γ antibody, 5 ng/ml TGF-β, 10 ng/ml IL-2 (PeproTech, 212-12). Cells were cultured in the presence of cytokines and antibodies for 72h and then harvested for FACS or Quantitative real-time PCR analysis.

### Quantitative real-time PCR

Total RNA was extracted using TRI reagent (Molecular Research Center, Inc., TR-118) and reverse transcribed into cDNA by mouse leukemia virus reverse transcriptase (Promega, M1708) and together with the Recombinant RNasin® ribonuclease inhibitor (Promega, N2518) to inhibit ribonuclease. Real-time PCR was performed in the Light Cycler quantitative PCR apparatus (Stratagene) with the 2×SYBR Green qPCR MasterMix (Bimake, B21702). β*-actin* was used as an internal standard gene, and the 2^-ΔΔCT^ method was utilized to quantitatively analyze the data. All qPCR primers used in this study are listed in the SI Appendix, Table S1.

### ELISA

Cell supernatants, mouse serum, and skin homogenate supernatants were collected. The levels of PAI-1 (R&D Systems, DY3828-05), IL-17A (Invitrogen, 88-7371-88), IL-6 (Invitrogen, 88-7064-88), and IL-23 (Invitrogen, 88-7230-88) were measured using a commercial enzyme-linked immunosorbent assay kit according to the instructions of the manufacturer. The absorbance was read in a microplate reader, and concentrations were calculated using a standard curve.

### Flow cytometry

Surface staining was performed using anti-mouse CD4 (Invitrogen, 25-0042-82) and anti-mouse CD8 (Invitrogen, 11-0081-85) antibodies. To assess cytokine expression, cells were stimulated with 50 ng/ml Phorbol-12-Myristate-13-Acetate (Sigma Aldrich, P1585), 750 ng/ml ionomycin (Sigma Aldrich, I3909), and 5 μg/ml brefeldin A (Thermo Fisher, 00-4506-5) for 4-6 h. Cells were then fixed and permeabilized using fixation/permeabilization solution reagent (Invitrogen, 88-8824-00), followed by intracellular cytokine staining with anti-mouse IFN-γ (BioLegend, 505810) and anti-mouse IL-17A antibodies (Invitrogen, 12-7177-81). For Treg cell staining, cells were fixed and permeabilized using a Foxp3 Fixation/Permeabilization Kit (Invitrogen, 00-5523-00) and stained with anti-mouse Foxp3 antibody (Invitrogen, 17-5773-82). Data were acquired on a BD FACS Verse System (BD Biosciences) and analyzed using FlowJo software (TreeStar).

### IMQ-induced psoriasis-like mouse model

Each mouse was topically treated daily with 62.5 mg of IMQ cream (5%) (Med-shine Pharma, H20030129) on the shaved back of 3.0 cm × 2.0 cm for 7 days to construct the psoriasis mouse model. The control mice were given the same dose of Vaseline cream. The Psoriasis Area and Severity Index (thickness, scaling, and erythema) parameters were used to assess disease severity in psoriasis mice. The severity of each parameter is graded as 0-4: 0=no clinical signs; 1=minor clinical signs; 2=moderate clinical signs; 3=significant clinical signs; 4=very significant clinical signs. On day 7, mice were euthanized, and tissue samples were collected for subsequent experiments.

### Histological analysis

Mouse skin tissues were fixed overnight with 4% paraformaldehyde (pH 7.2), embedded in paraffin, and sliced into 5 μM sections. To assess skin thickness and inflammation, skin sections were stained with hematoxylin and eosin. Images were captured using an Olympus IX51 inverted fluorescent microscope, and quantitative image analysis was performed using ImagePro.

### Statistical analysis

Data were presented as mean ± SEM and analyzed using GraphPad Prism 8. For comparisons of variables between two groups, the unpaired Student’s t-test was used. One-way ANOVA was used to compare more than two groups. Two-way ANOVA was used to compare the differences in clinical scores of psoriasis model mice between groups. P<0.05 was considered statistically significant.

### Data, Materials, and Software Availability

The material and data used to support the current study’s findings are available from the corresponding author upon reasonable request. The publicly available data sets used in this study are indicated accordingly.

## Acknowledgments

We gratefully acknowledge the staff of the College of Life Sciences and Technology at Tongji University. We thank the members of the Du lab for their critical comments and discussions. This work was supported by funds from the National Natural Science Foundation of China (32070768, 32270754).

## Author contributions

K.S., L.X., C.L., and C.D. designed research; K.S., L.X., C.W., J.L., and G.L. performed research; S.H., J.L., and C.D. contributed analytic tools; K.S., L.X., and C.D. analyzed data; C.L. and C.D. supervised the entire study; K.S., C.D., and C.L. wrote the paper.

## Supporting Information

**Fig. S1.**
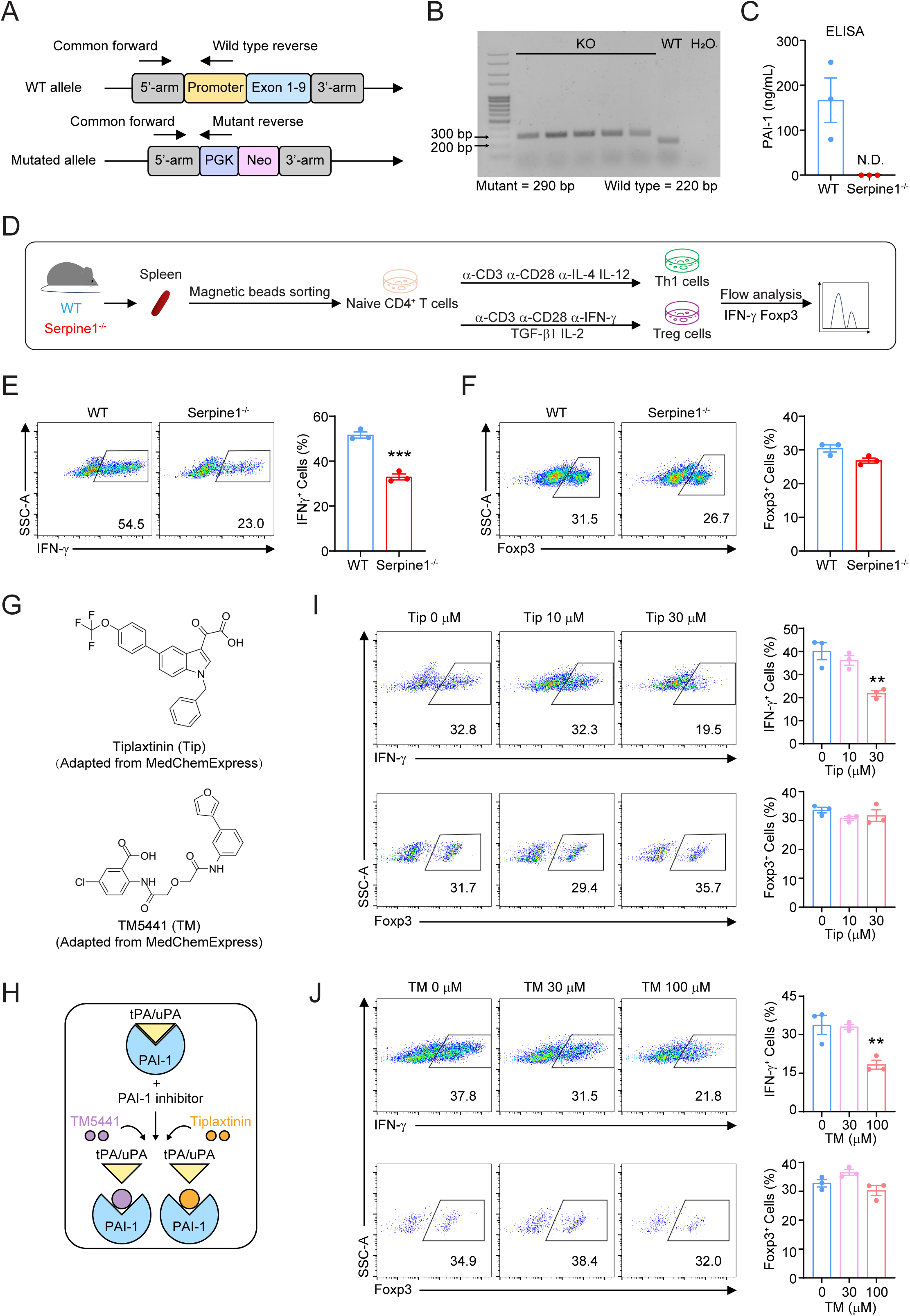
Serpine1 deficiency affects Th1 but not Treg cell differentiation. (A) Location of primers for Serpine1^-/-^ mouse identification. (B) PCR detection of tail DNA to determine the genotype of progeny. (C) ELISA assay of PAI-1 expression levels in serum of WT and Serpine1^-/-^ mice. (D) Schematic diagram of Th1 and Treg cell differentiation *in vitro*. (E) Flow cytometry detection of Serpine1^-/-^ and WT Th1 cells (IFN-γ^+^) differentiated *in vitro*. Representative flow cytometry images (left). IFN-γ^+^ cell frequency (right). (F) Flow cytometry detection of Serpine1^-/-^ and WT Treg cells (Foxp3^+^) differentiated *in vitro*. Representative flow cytometry images (left). Foxp3^+^ cell frequency (right). (G) Schematic structure of the PAI-1 inhibitors Tiplaxtinin and TM5441. (H) Schematic diagram of the mechanisms of the PAI-1 inhibitors Tiplaxtinin and TM5441. (I) Flow cytometry detection of the differentiation ratios of Th1 cells (IFN-γ^+^) and Treg (Foxp3^+^) cells differentiated *in vitro* in the presence of 0, 10, and 30 μM Tiplaxtinin. Representative flow cytometry images (left). Cell frequencies (right). (J) Flow cytometry detection of the differentiation ratios of Th1 cells (IFN-γ^+^) and Treg (Foxp3^+^) cells differentiated *in vitro* in the presence of 0, 30, and 100 μM TM5441. Representative flow cytometry images (left). Cell frequencies (right). Data are presented as mean ± SEM. Statistical analysis was carried out by unpaired Student’s t-test (E and F) or one-way ANOVA test (I and J). \*\**P*<0.01, \*\*\**P*<0.001.

**Fig. S2.**
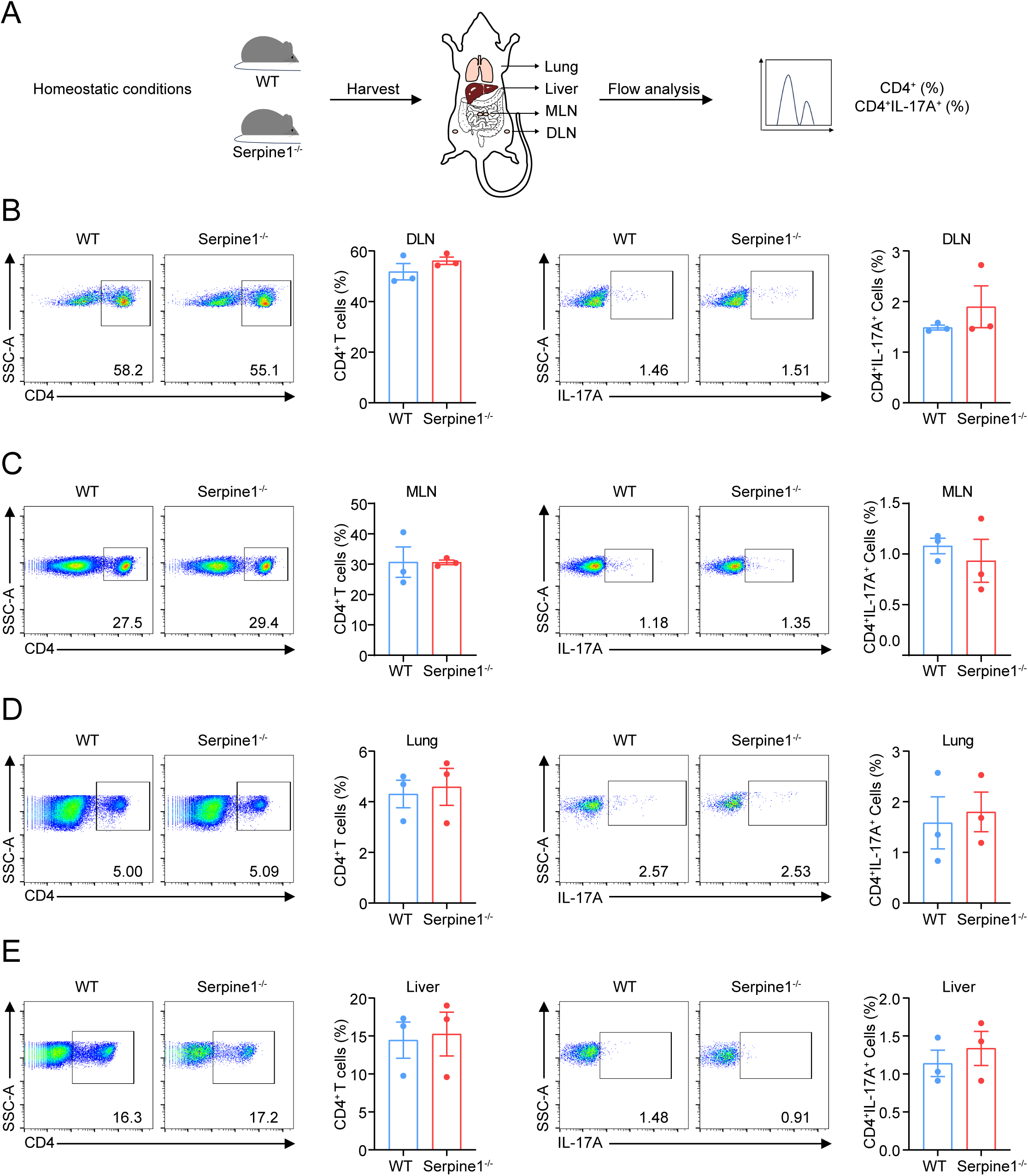
Serpine1 deletion at steady state has no effect on Th17 cells in different tissues. (A) Flow cytometry detection of CD4 cell (CD4^+^) and Th17 cell (CD4^+^IL-17A^+^) ratios in the DLN, MLN, lungs, and livers of Serpine1^-/-^ and WT mice. (B) Flow cytometry detection of the ratio of CD4 cells (CD4^+^) and Th17 cells (CD4^+^IL-17A^+^) in the DLN of Serpine1^-/-^ and WT mice. Representative flow cytometry images (left). Cell frequencies (right). (C) Flow cytometry detection of the ratio of CD4 cells (CD4^+^) and Th17 cells (CD4^+^IL-17A^+^) in the MLN of Serpine1^-/-^ and WT mice. Representative flow cytometry images (left). Cell frequencies (right). (D) Flow cytometry detection of the ratio of CD4 cells (CD4^+^) and Th17 cells (CD4^+^IL-17A^+^) in the lung of Serpine1^-/-^ and WT mice. Representative flow cytometry images (left). Cell frequencies (right). (E) Flow cytometry detection of the ratio of CD4 cells (CD4^+^) and Th17 cells (CD4^+^IL-17A^+^) in the liver of Serpine1^-/-^ and WT mice. Representative flow cytometry images (left). Cell frequencies (right). Data are presented as mean ± SEM. Statistical analysis was carried out using an unpaired Student’s t-test (B-E).

**Fig. S3.**
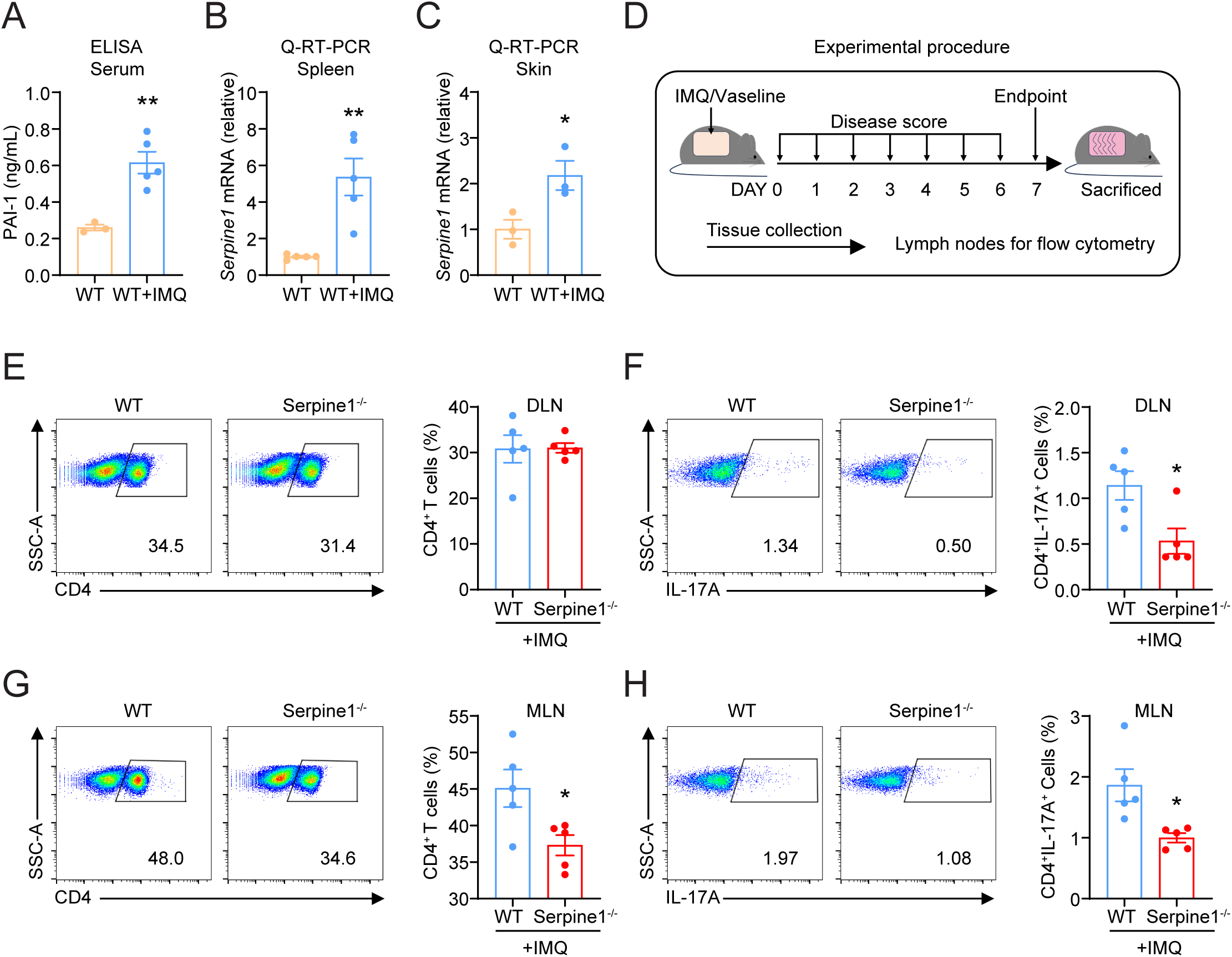
Seprine1 deficiency affects Th17 cells in the lymph nodes of psoriatic mice. (A) Detection of PAI-1 expression level in mouse serum by ELISA. (B-C) Detection of Serpine1 mRNA levels in mouse (B)spleen and (C)skin by q-RT-PCR. (D) Schematic diagram of flow cytometry detection of CD4 and Th17 cells in the lymph nodes of mice with psoriasis. (E and G) Flow cytometry to detect the proportion of CD4 cells (CD4^+^) in the (E) DLN or (G) MLN of psoriatic mice. Representative flow cytometry images (left). Cell frequency (right). (F and H) Flow cytometry to detect the proportion of Th17 cells (CD4^+^IL-17A^+^) in the (F) DLN or (H) MLN of psoriatic mice. Representative flow cytometry images (left). Cell frequency (right). Data are presented as mean ± SEM. Statistical analysis was carried out using an unpaired Student’s t-test (H-N). \**P*<0.05, \*\**P*<0.01.

**Fig. S4.**
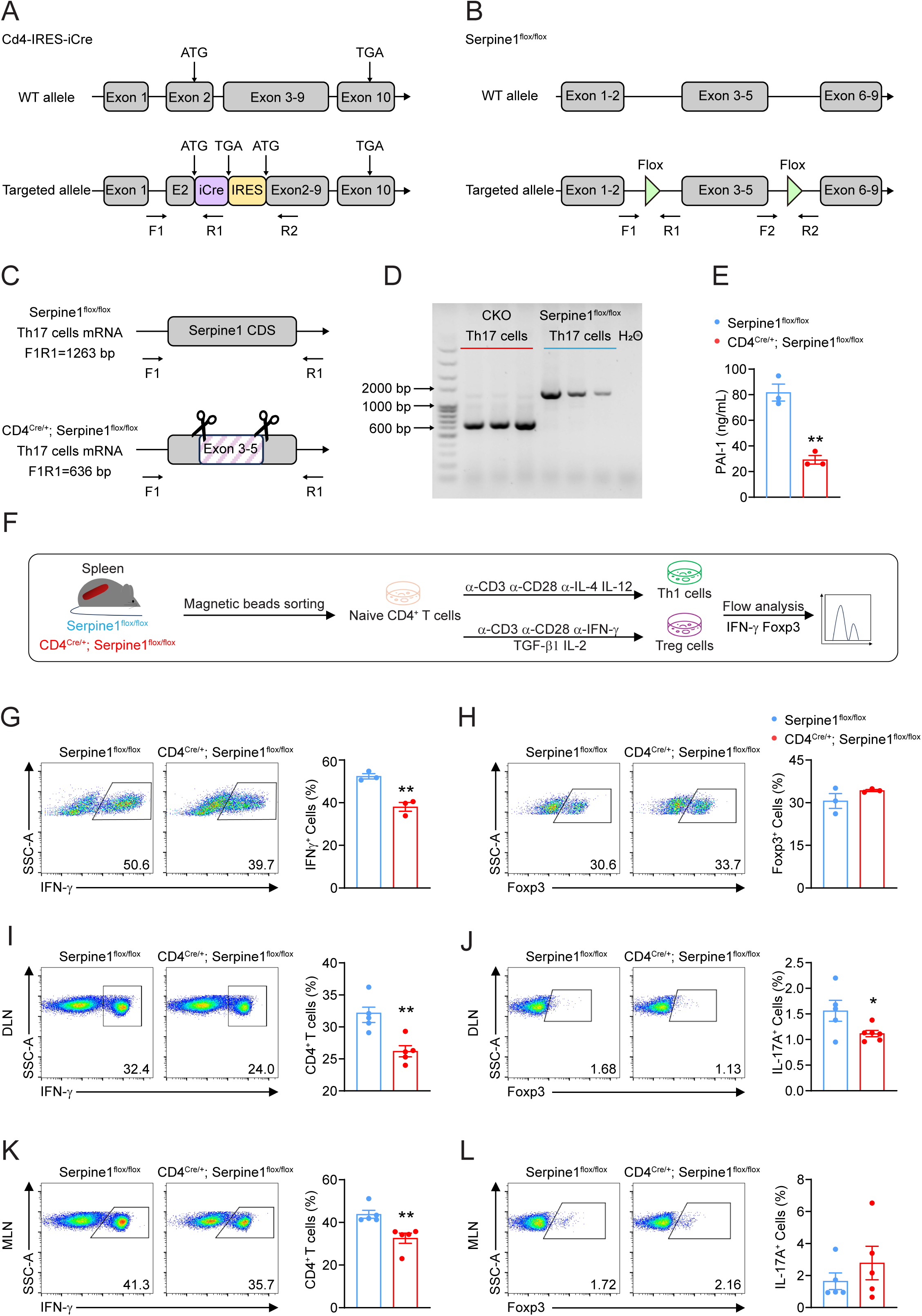
Generation of CD4-specific serpine1 cKO mice. (A) The schematic diagram of Cd4-IRES-iCre mouse generation. (B) The schematic diagram of Serpine1^flox/flox^ mouse generation. (C-E) *In vitro* differentiated Serpine1^flox/flox^ and CD4^Cre/+^; Serpine1^flox/flox^ Th17 cells and supernatants were collected. RNA was extracted from Th17 cells and reverse transcribed into cDNA. The PCR product of Serpine1 in Serpine1^flox/flox^ mice is 1263 bp, whereas the cDNA region in cKO mice is 636 bp in length due to the absence of exons 3-5 (627 bp). (D) Presentative PCR detection of cDNA length in Serpine1^flox/flox^ and CD4^Cre/+^; Serpine1^flox/flox^ Th17 cells. (E) Detection of PAI-1 in the supernatants of Th17 cells differentiated *in vitro* from Serpine1^flox/flox^ and CD4^Cre/+^; Serpine1^flox/flox^. (F) Schematic diagram of Th1 and Treg cell differentiation *in vitro*. (G) Flow cytometry detection of Serpine1^flox/flox^ and CD4^Cre/+^; Serpine1^flox/flox^ Th1 cells (IFN-γ^+^) from *in vitro* differentiation. Representative flow cytometry images (left). IFN-γ^+^ cell frequency (right). (H) Flow cytometry detection of Serpine1^flox/flox^ and CD4^Cre/+^; Serpine1^flox/flox^ Treg cells (Foxp3^+^) differentiated *in vitro*. Representative flow cytometry images (left). Foxp3^+^ cell frequency (right). (I and K) Flow cytometry to detect the proportion of CD4 cells (CD4^+^) in the (I) DLN or (K) MLN of psoriatic mice. Representative flow cytometry images (left). Cell frequency (right). (J and L) Flow cytometry to detect the proportion of Th17 cells (CD4^+^IL-17A^+^) in the (J) DLN or (L) MLN of psoriatic mice. Representative flow cytometry images (left). Cell frequency (right). Data are presented as mean ± SEM. Statistical analysis was carried out using an unpaired Student’s t-test (H-N). \**P*<0.05, \*\**P*<0.01.

---

**Table S1.**
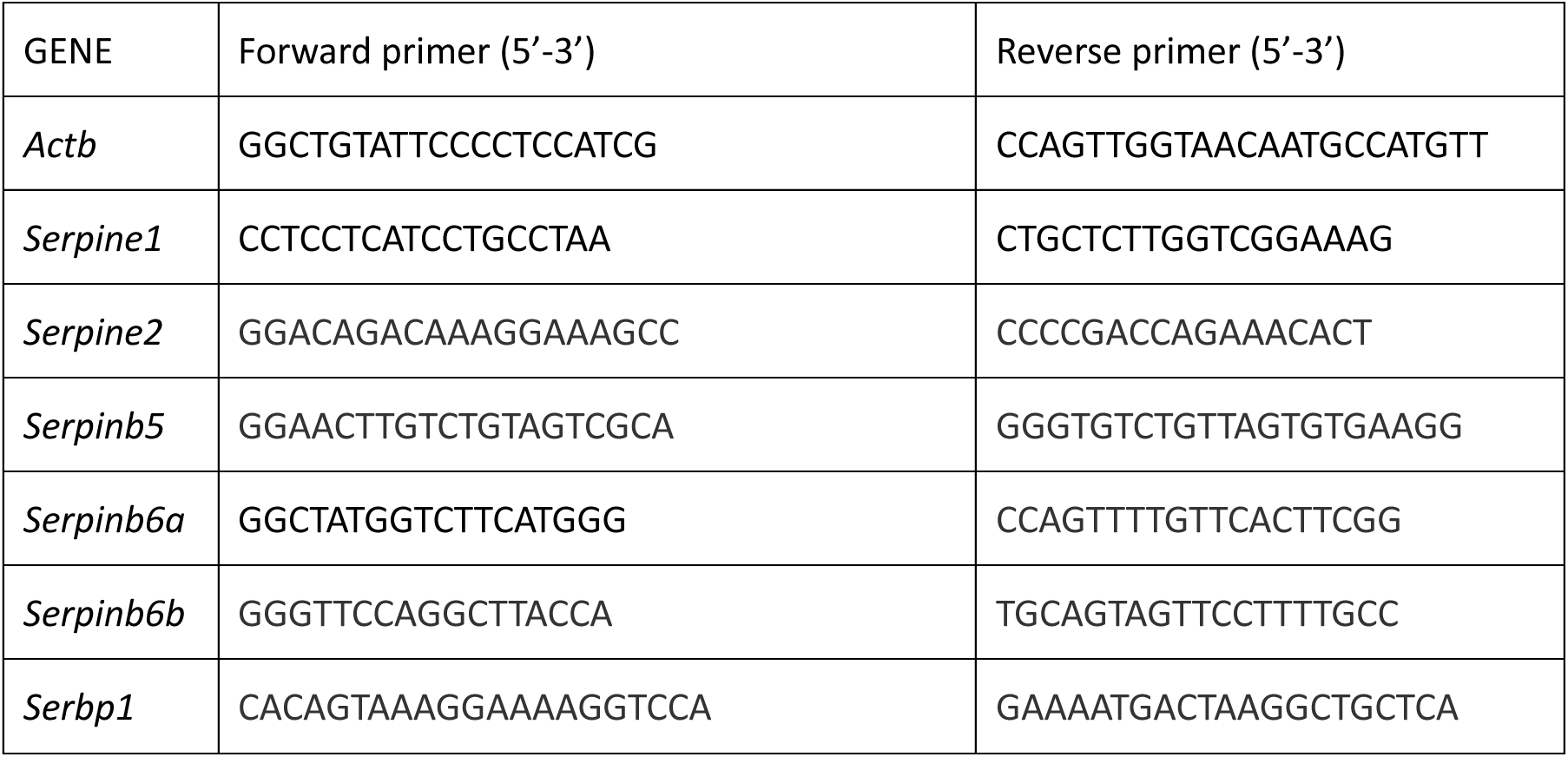
Primer sequences used for RT-PCR analysis.

